# Human origin and migration deciphered from a novel genomic footprints of mitochondrial sequences

**DOI:** 10.1101/848341

**Authors:** Aritra Mahapatra, Jayanta Mukherjee

**Affiliations:** Department of Computer Science and Engineering, Indian Institute of Technology, Kharagpur, India, 721302

**Keywords:** Human migration, modern humans, mitochondrial genome, graphical representation, graphical footprint, drift

## Abstract

The origin of modern human and their migration across the world is one of the most debated topics for the decades. There exist two different hypotheses, recent African origin and multi-regional evolution, based on the genomic studies, haplogroups, archaeological records, cultural behaviors, palaeontology studies, etc. Various studies placed the modern humans in a phylogenetic tree to depict the relationships among them. The debate for determining those regions of Africa which witnessed the first origin of humans still exists. The conflicts between the results obtained from the molecular data and the archaeological and palaeontological reports still exist. We adopt a novel genomic feature derived from the whole mitochondrial sequence, and using a novel distance function the phylogenetic trees are constructed based on the feature which provide a new insight on human migration. We propose a new method to derive the bootstrap replica from the genome sequences by considering the genetic variance to demonstrate the robustness of the obtained trees. The results derived from the genomic feature are more consistent with the archaeological findings based on the time of origin of different communities. We find that west and central African communities are placed at the basal point with a very high bootstrap score. This study roughly estimates the existence of the archaic human at 800-900 kilo years ago and presence of human in Africa at 600-700 kilo years ago. This supports the presence of an ancestor in the west and central Africa much earlier than that of the fossils identified.

## 1 Introduction

The origin of the anatomically modern humans (AMH) and their migration into the world is being studied extensively. The multiple genetic admixtures due to migration, intermarriage, slavery, human trafficking etc. played an important role to make the human history very complex (Duda and 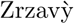, 2016; Mellars, 2006). Apart from that, different assumptions such as gene flow (Shriner et al., 2016), make the study of human evolution and migration more complex (Bae et al., 2017).

Broadly, there exist two hypotheses of human evolution and migration. The “recent African origin” or “replacement” hypothesis (Chan et al., 2019; Cann et al., 1987; Vigilant et al., 1991) states that the already existing anatomically modern humans subsequently replace the ancient humans and spread all over the world (Liu et al., 2006; Stringer, 2003; Wolpoff et al., 2000). There are various methods incorporated to describe the history of the humans. Several genomic studies (Mondal et al., 2019; Browning et al., 2018; Mondal et al., 2016) supported the “replacement” hypothesis. This hypothesis proposes that there is no or very less admixture or genetic mixing between AMH and the archaic humans throughout the world. But this hypothesis is negated by some recent studies (Mafessoni, 2019; Prüfer et al., 2014). Alternatively, the “multi-regional” hypothesis (Cartmill and Smith, 2009) states that the evolution of the archaic human occurred in different parts of the world. This evolution ended up with the AMH. Several linguistic analysis (Baker et al., 2017; Balter, 2019), and cultural analysis (Bhugra and Becker, 2005; Mesoudi, 2018) support the “multi-regional” hypothesis.

Though various genomic studies were conducted for searching the origin of humans, most of the studies focused on the phylogenetic tree derived from alignment based methods (Mondal et al., 2019; Jinam et al., 2017; Töpf et al., 2005; Basu et al., 2003), SNP genotyping (Juyal et al., 2014; Moorjani et al., 2013), or supertree based approach (Duda and 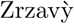, 2016). Since then, the conflicts between the genomic studies and the archaeological and palaeontological studies exist in the question of the origin of humans (Callaway, 2018). As a result many archaeologists think the ancient DNA as the “devil’s work” (Callaway, 2018). Though the origin of the human in the eastern and southern parts of Africa has long been accepted (Sahle et al., 2018; Behar et al., 2008; Chan et al., 2015, 2019), a few recent studies documented the presence of humans in the western part of Africa much earlier than the other fossils identified till date (Stringer and Galway-Witham, 2017; Durvasula and Sankararaman, 2020). In this work, we consider the mitochondrial genome (mtDNA) for studying the origin of humans. The mtDNA is haploid, inherited maternally (Hurles and Jobling, 2001), and recombination is very rare event in it (Eyre-Walker and Awadalla, 2001). So the changes of mtDNA sequence occur mainly due to mutations. Our study shows a consistency with the archaeological and palaeontological studies based on the age of origin of human in Africa. Moreover, this study also supports the recently published studies on the origin of human in Africa.

In our present study, we use a novel feature representation of the whole mitochondrial genome (mtDNA) based on a graphical encoding method, named GRAFree (Mahapatra and Mukherjee, 2019). We convert a whole mtDNA sequence into a fixed dimensional feature vector. This feature vector is used for studying the phylogeny of human representatives of different regions of this modern world. It has been hypothesized that hereditary relationship among humans of different regions is reflected by their closeness in genomic signatures, in this case, in mitochondrial genomes (Langille et al., 2010). Discovering this relationship and building a phylogenetic hierarchy among representatives of different parts of this world is the key motivation behind the mathematical representation of a genome. We plot the mtDNAs in a two dimensional coordinate plane by considering successive shifts in *x* and *y* coordinates using three different encodings of DNA alphabets (see Methods). The 2-D distribution of points are represented by a 5-D vector, formed by the mean, eigen values, and the direction of the eigen vector corresponding to the major eigen value. We adopt a distance function to compute the pair-wise distances using the feature vectors. This comparative study finally gives us a distance matrix. We apply a hierarchical clustering method, UPGMA (Sneath and Sokal, 1973), to derive the phylogenetic tree. We consider three different encodings of shifts of coordinates corresponding to the nucleotides in the genomic sequence, and generate three phylogenetic trees independently. Finally, we combine the trees using a quartet based supertree construction method, ASTRAL (v4.10) (Sayyari and Mirarab, 2016). We believe that, this is the first study of human origin based on the graphical representation of genome.

We apply this method on a large set of mitochondrial DNA sequences of humans. In this study, we consider 369 mitochondrial genomes from more than 120 countries covering six continents (except Antarctica). The data are sequenced and published by various researchers (Friedlaender et al., 2005; Ingman et al., 2000; Roostalu et al., 2006; Hill et al., 2006; Achilli et al., 2005; Olivieri et al., 2006; Maca-Meyer et al., 2001; Starikovskaya et al., 2005; Behar et al., 2006; Ingman and Gyllensten, 2003, 2007). They are accumulated and stored in a database server named *mtDB* (Ingman and Gyllensten, 2006). Apart from that we include six outgroup species in our study (see **Supplementary S1**). We found that GRAFree distinguishes the outgroups from the humans with very high bootstrap supports. The output shows that the time of appearance of human in Africa is consistent with the archaeological and palaeontological studies. We found that a significantly large population from the western and central parts of Africa are placed at a distinct clade as compared to the rest of the humans with a very high bootstrap support indicating a divergence of the genomic signatures of the western and central African population from the rest of the humans. This phenomena also indicates an early origin of human from these parts of Africa to the world. This period is roughly estimated to be 600-700 kilo years ago (kya). This may be earlier than the period presently being considered as the period of traceable human origin to these regions, which is between 230-300 kya (Andrews and Johnson, 2019; Smith and Ahern, 2013). This study also indicates presence of lineages of migrated humans to this part of world which took place much ahead of 65 kya, what is considered to be the present time line for the out of Africa hypothesis (Veeramah and Hammer, 2014). Our finding is consistent with archaeological findings and discoveries of ancient human skeletons, which predate the estimated periods from analysis of different mtDNA and Y-Chromosomal haplogroups of human population. It provides a new insight to the “out of Africa” hypothesis. In addition, this study also indicates the ancestor of the African populations of western and the central parts appeared much earlier than the period being documented as the origin of Neanderthals and Denisovans (Veeramah and Hammer, 2014; Green et al., 2010; Mendez et al., 2012).

## 2 Materials and methods

We consider three sets of structural groups of nucleotides (purine, pyrimidine), (amino, keto), and (weak H-bond, strong H-bond) separately for representing DNA by a sequence of points in a 2-D integral coordinate space. This point set is called **Graphical Foot Print (GFP)** of a DNA sequence. We adopt a technique for extracting features from GFPs and use them for constructing phylogenetic trees (Mahapatra and Mukherjee, 2019). As there are three different types of numerical representation of nucleotides, we could form three different hypotheses for the phylogeny. Finally, we combine the trees generated from these three hypotheses by applying a tree merging algorithm called ASTRAL (Mirarab et al., 2014; Sayyari and Mirarab, 2016).

### 2.1 Feature space

#### Definition 1 Graphical Foot Print (GFP)

*Let* 𝒮 *be a DNA sequence, such that*, 𝒮 ∈ ∑^+^, ∑ = {*A, T, G, C*}. *For each combination of Purine (R)/Pyrimidine (Y), Amino (M)/Keto (K), and Strong H-bond (S)/Weak H-bond (W), the GFP of* 𝒮, *ϕ*(𝒮), *is the locus of 2-D points in an integral coordinate space, such that* (*x*_*i*_, *y*_*i*_) *is the coordinate of the alphabet s*_*i*_, ∀ *s*_*i*_ ∈ 𝒮, *for i* = 1, 2, *…, n, and x*_0_ = *y*_0_ = 0.

*Case-1: for Purine/Pyrimidine*

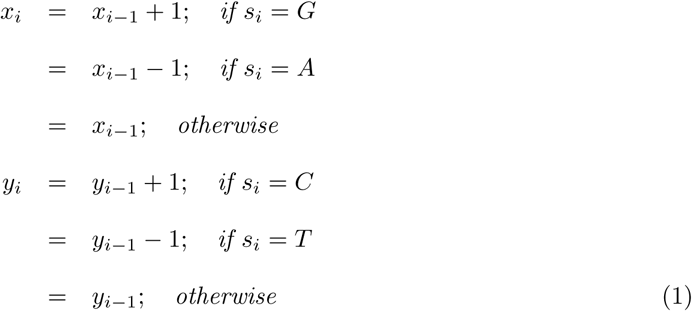

*Case-2: for Strong H-bond/Weak H-bond*

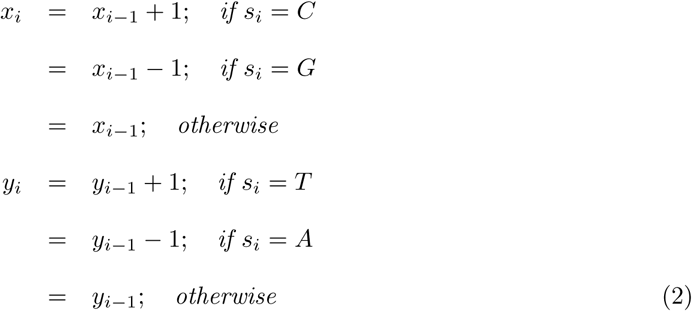

*Case-3: for Amino/Keto*

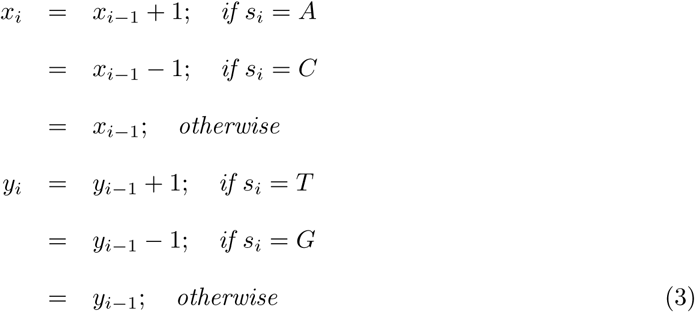

We denote GFPs of Case-1, Case-2, and Case-3, as GFP-RY (Φ_*RY*_), GFP-SW (Φ_*SW*_), and GFP-MK (Φ_*MK*_), respectively.

#### Definition 2 Drift of GFP

*Let* 𝒮 *be a DNA sequence and s*_*i*_ *be the alphabet (s*_*i*_ ∈ {*A, T, G, C*}*) at the i*^*th*^ *position of* 𝒮. *Let ϕ*_*i*_(𝒮) *denote the corresponding (x*_*i*_,*y*_*i*_*) the coordinate of s*_*i*_ *in* Φ(𝒮).

*Then for length L, drift at the i*^*th*^ *position is defined as*,

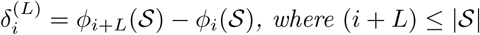

*Considering the drifts for every i*^*th*^ *location of the whole sequence, the sequence of drifts is denoted by*

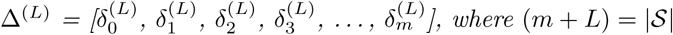

For GFP-RY (refer to Definition 1), an element (*x*_*i*_, *y*_*i*_) in 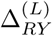 provides excess numbers of *G* from *A* and *C* from *T* in segment of length *L* starting from the *i*^*th*^ location, respectively. Similarly, in GFP-SW, they are the excess numbers of *C* from *G* and *T* from *A* (represents as 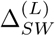), and in GFP-MK, they correspond to the excess numbers of *A* from *C* and *T* from *G* (represents as 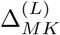), respectively.

We also call the elements of Δ^(*L*)^ as points, as they can be plotted on a 2-D coordinate system. We call this plot as the scatter plot of the drift sequence. Similarly, we get a scatter plot of a GFP. It has been observed that in many cases the scatter plots of Δ have similar structure for closely spaced species mentioned in literature. It can also be observed that differences between two GFPs get reflected in their respective drifts.

We represent spatial distribution of these points of Δ by an elliptical model using a five dimensional feature descriptor: (*µ*_*x*_, *µ*_*y*_, Λ, *λ, θ*), where (*µ*_*x*_, *µ*_*y*_) is the center of the coordinates, Λ and *λ* are major and minor eigen values of the covariance matrix, and *θ* is the angle formed by the eigen vector corresponding to Λ with respect to the *x*-axis. We make ℱ number of non overlapping equal length fragments from Δ and represent each fragment using the five dimensional feature descriptor.

### 2.2 Distance function

For two sequences 𝒫 and 𝒬 with the feature descriptors of *i*^*th*^ fragments 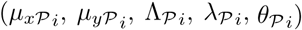 and 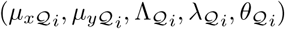, where *i* ≤ ℱ, we use the following distance function between them,

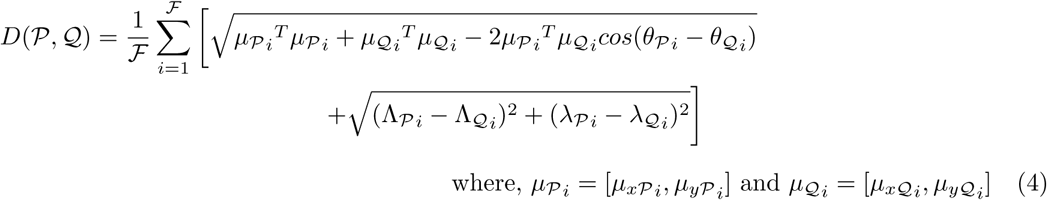

It can be shown that the above distance function is a metric.

## 3 Results

### 3.1 Computation of GFP and Drift sequence

In this study, we consider three sets of structural groups of nucleotides (purine, pyrimidine), (amino, keto), and (weak H-bond, strong H-bond) separately for representing DNA by a sequence of points in a 2*D* integral coordinate space. We derive the GFP using these three encodings. We consider them as *GFP*_*RY*_, *GFP*_*SW*_, and *GFP*_*MK*_, respectively. The drift sequence of a GFP is the coordinate difference between those pair of points which are spanned by *L* number of points in between. Our proposed method, GRAFree, depends on two parameters such as the size of the window (*L*) and number of fragments of the sequence of drifts (ℱ). Each fragment is represented by a 5*D* vector as mentioned above and concatenation of ℱ vectors provide a feature vector of dimension 5ℱ.

Now, what is the optimal value of *L* and ℱ which we should consider for the input dataset. For this, we compute the Shannon entropy (Shannon, 2001) of the drift sequence of each GFP for different values of *L* (starting from 50 to 2000 with the difference of 50). It is observed in Fig. 1 that for all of the GFPs initially by increasing the value of *L*, the entropy increases till a certain limit. This incident represents that by increasing the value of *L*, the number of distinct points of the sequence of the drifts also increases. After a certain value of *L*, e.g, at *L* = 550, the entropy of all mtDNA becomes stabilized (changes *<* 1%). By increasing the value of *L* after that does not change the entropy significantly. It is observed that for *L* ≥ 550, the Δ^(*L*)^ contains significantly large number of distinct point coordinates than that of *L <* 550. Hence, we choose the value of *L* as 550.

**Figure 1:**
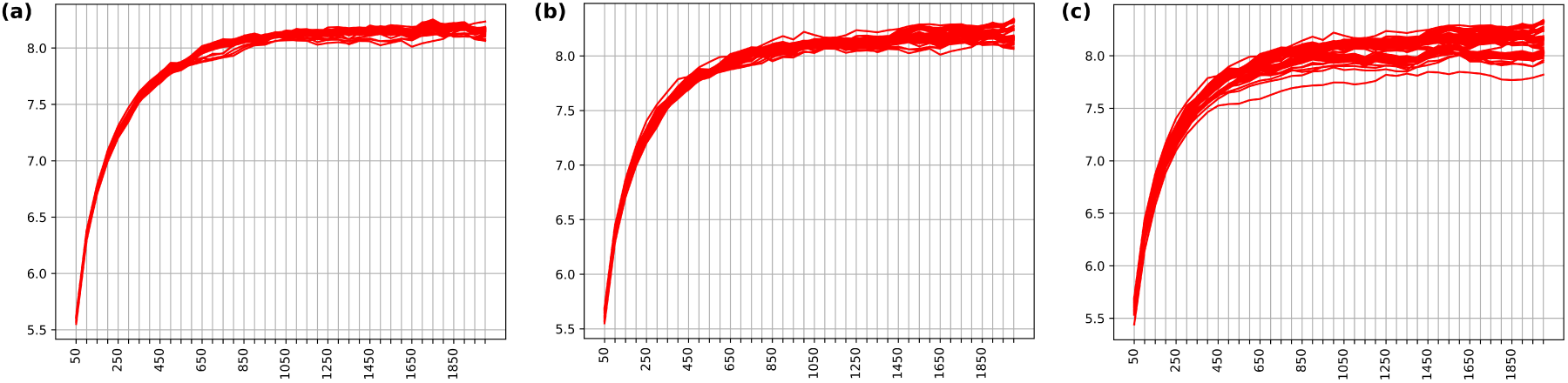
Shannon entropy of (a) *GFP*_*RY*_, (b) *GFP*_*SW*_, and (c) *GFP*_*MK*_ of different human mtDNA for all the values of *L* from 50 to 2000.

For getting more precise signature of a drift sequence, we make 15 non-overlapping partitions of the drift sequence. Hence, in this study, we consider ℱ as 15. Each fragment is represented by the 5*D* feature vector which consists of the mean of the point coordinates of the corresponding fragment, eigen (both major and minor) values, and the angle formed by the eigen vector corresponding to the major eigen value with respect to the *x*-axis. Hence, a complete drift is represented by 75 dimensional feature vector.

We compute the Graphical Foot Print (GFP or *ϕ*) of each mtDNA of human taken from different parts of the world. For *L* = 550, we also compute the sequence of the drifts of the corresponding GFP. It is observed that the species which are closely related, have similar patterns in both of their GFP and drift sequence. Typical examples of drift sequences of humans from different regions of the world and the outgroup species, such as baboon, gorilla, and chimpanzee are shown in Fig. 2.

**Figure 2:**
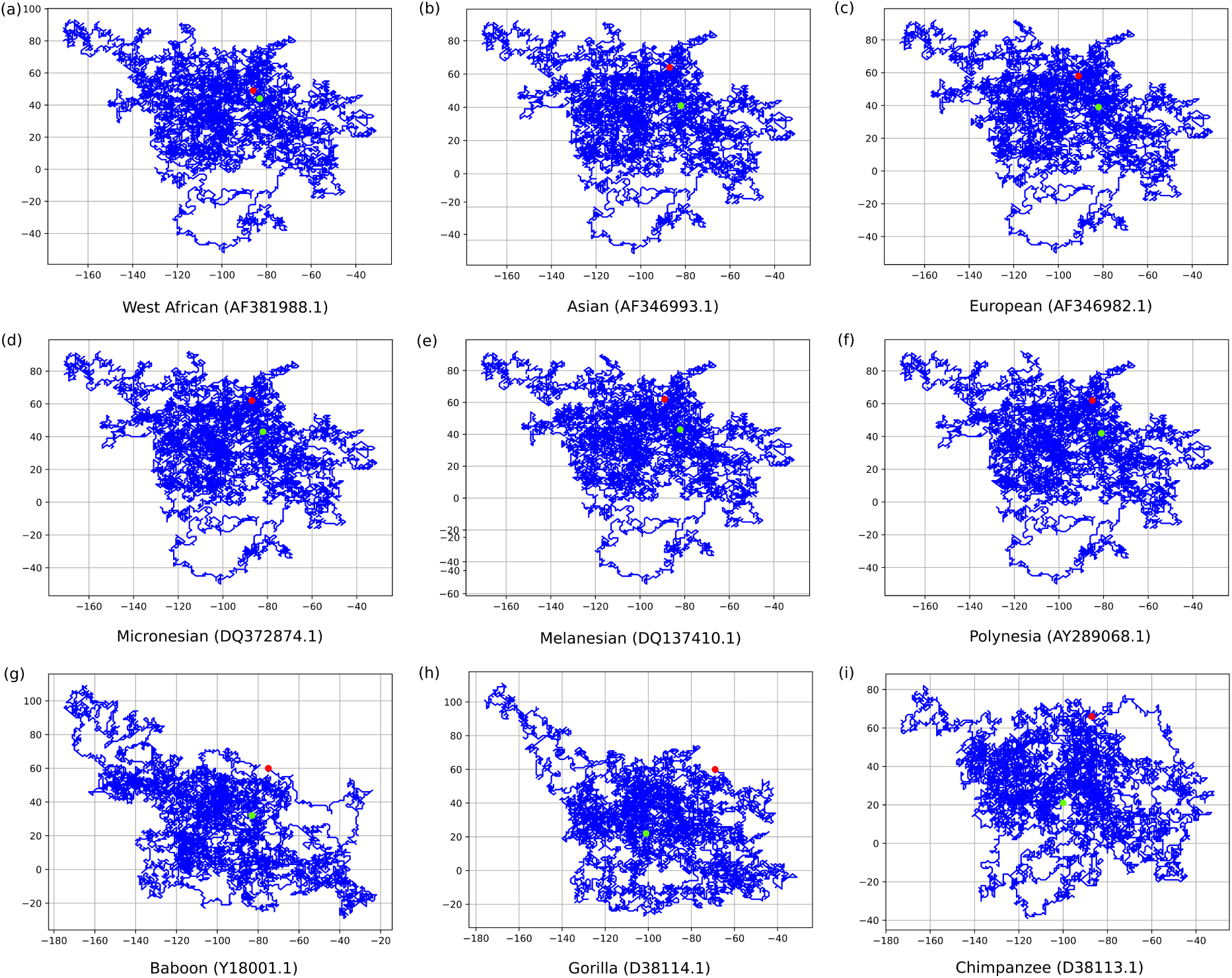
The sequence of drifts of different sample, mentioned under each graph, for the case when we consider the Purine and Pyrimidine as 2*D* coordinates space. The accession numbers of the corresponding samples are mentioned within the bracket. The start and end positions of the sequence of drifts are denoted by the green and red dots, respectively. The drift sequences of the Baboon, Gorilla, and Chimpanzee (g,h,i) are much different from the drift sequences of human (a-f). The drift sequences of the mtDNA of the population of the western Africa (a) are slightly different from the drift sequences of the other humans (b-f). The differences are visible to the naked eye at the starting (green dot) and end points (red dot) of the drift sequences.

### 3.2 Deriving trees

We propose a novel distance function to compute the pair-wise distances. Our proposed distance function is a metric. We apply a hierarchical clustering method, Unweighted Pair Group Method with Arithmetic mean (UPGMA) (Sneath and Sokal, 1973), to form the phylogenetic tree from the distance matrix.

We apply this same technique on three different cases such as *GFP*_*RY*_, *GFP*_*SW*_, and *GFP*_*MK*_, separately. Thus we derive three different phylogenetic trees by following three different hypotheses. These trees are provided in the **Supplementary S2 Fig. 1-3 in Section 1**. There are various methods to combine multiple trees and to form a supertree. We apply a quartet based method, ASTRAL (Sayyari and Mirarab, 2016), to combine three trees. Please refer to Fig. 3 and also refer to the **Supplementary S2 Fig. 4 in Section 1**.

**Figure 3:**
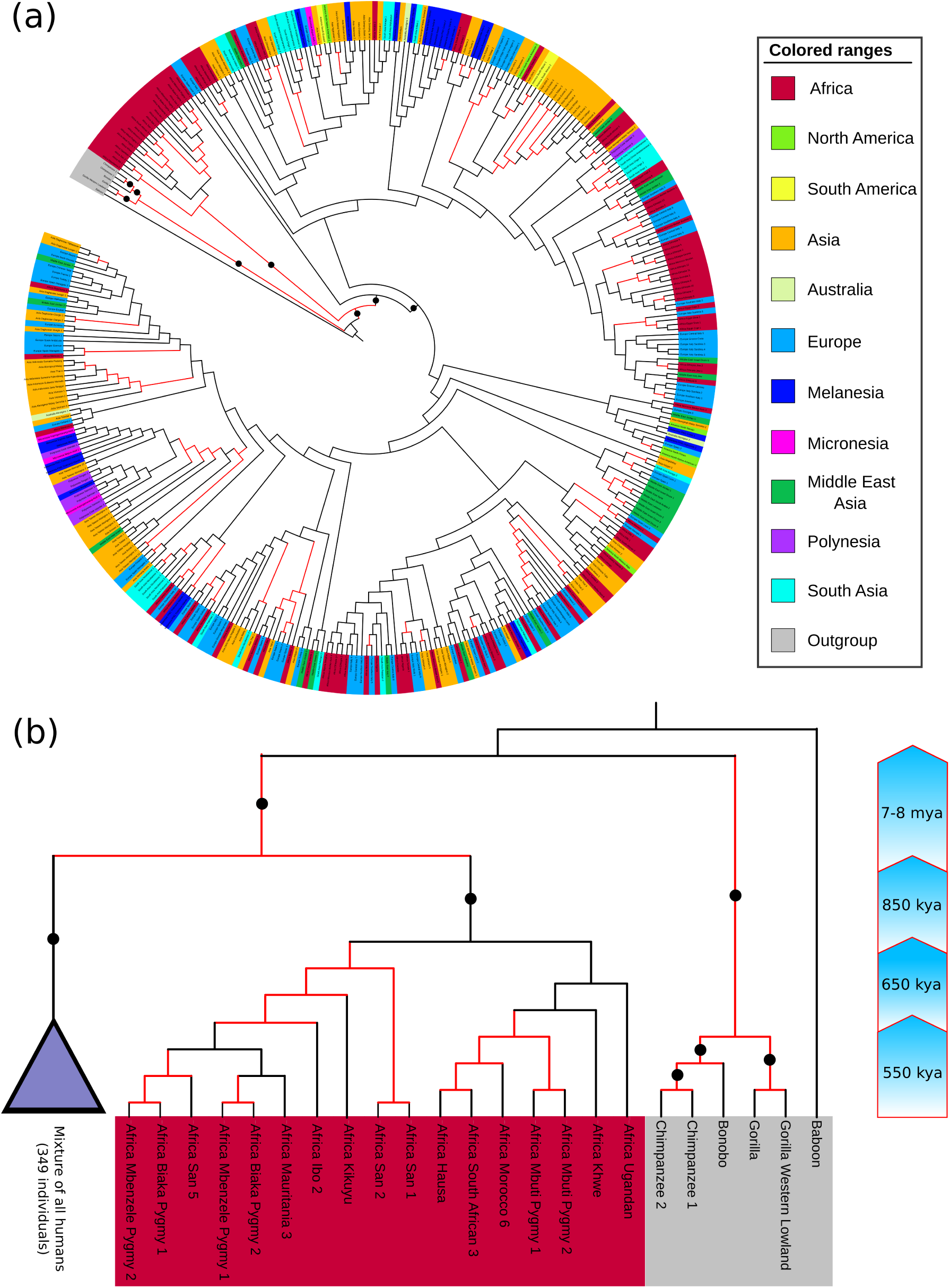
The derived trees using GRAFree. The black dots on the edges denote a high bootstrap support (≥ 75%) of the corresponding clade. The edge marked as red color denotes that the robustness of the clade with respect to the hyper parameters is greater than 80%. (a) Tree derived after merging all three trees using ASTRAL. (b) The summarization of the tree having bootstrap support ≥ 75%.

### 3.3 Bootstrapping

The primary motivation of bootstrapping is to generate the population from a single genome. It is observed that the average intraspecies variation in mitochondrial genome of *Homo sapiens* is within 1.2% (Castro et al., 1998). But in this study, we are trying to derive the relationships between different communities of human. Hence, we consider the average genetic variation of mtDNA for a particular community as 0.3% (Jarczak et al., 2019). Here we propose a bootstrapping technique which considers the genetic variance of a sample space within 0.3%. For that we apply insertion, deletion, and mutation at each location with an equal probabilty of 0.1% and consider an unbiased selection of the nucleotides at each location for the insertion and the mutaion. We generate 100 replicas using this proposed bootstrapping method and construct trees from each set of sequences using GRAFree method by setting values of *L* and ℱ as discussed in the previous section. Felsenstein’s bootstrapping method (Felsenstein, 1985) assesses the robustness of phylogenetic trees using the presence and absence of clades. For the large scale genomics Felsenstein’s bootstrap is not efficient to sum up the replicas. For the hundred of species this method is inclined to produce low bootstrap support (Lemoine et al., 2018). So here we apply a modification of Felsenstein’s bootstrapping, where the presence of a clade is quantified using the transfer distance proposed in (Lemoine et al., 2018). The transfer distance (Charon et al., 2006) is the minimum number of changes required to transform one partition to another. We compute the occurrence of each clade using the tool BOOSTER (Lemoine et al., 2018). The clades having a high bootstrap support (≥ 75%) are denoted as the black dot on the edge in Fig. 3 and also refer to the **Supplementary S2 Fig. 1-4 in Section 1**.

### 3.4 Robustness against variation of hyper parameters

Here we test the consistency of the results for different values of *L* and ℱ. For that we generate a large number of trees by considering the values of *L* from 500 to 1000 and ℱ from 5 to 50. We consider the tree derived for *L* = 550 and ℱ = 15 as the reference tree. It is observed that most of the clades of the reference tree is consistent (occurrences more than 80%) for different parameter values (Fig. 3) and also refer to the **Supplementary S2 Fig. 1-4 in Section 1**.

### 3.5 Estimation of time of origin of clades

We applied UPGMA technique to get the edge lengths of the derived trees. Most of the studies proposed that the speciation of chimpanzee and human occurred 7-8 million years ago (mya) (Langergraber et al., 2012; Patterson et al., 2006; Reich, 2018). To calibrate the distances of the derived trees we consider the speciation age of chimpanzee and human and derived the age of the origin of different groups of humans. We have used this calibration to estimate time of origins of clades which have strong bootstrap supports. It is observed that the most recent common ancestor of all the humans originated about 850 kya, while the African clade originated 650 kya. The derived trees also indicate that the “Out-of-Africa” migration appeared about 550 kya (Fig. 3). The trees with the edge lengths are provided in the **Supplementary S3**.

## 4 Discussion

We develop a new approach, GRAFree, to derive the phylogenetic relationships among a large number of mitochondrial genome of AMH of different regions of this world. We also include six outgroup species (such as other than human) in our study. The proposed approach is quite different from mitochondrial haplogroup based approaches. This method is effective in capturing broad distinct clades. This is evident from successful separation of the outgroup species from human in our study. However, this method identifies close relationship between chimpanzee and gorilla than human. It is also observed that the proposed distance function is more robust than the Euclidean distance function for the selected feature vector.

It is a well accepted hypothesis that the ancient human originated in Africa. The southern and eastern Africa is long been considered as the region where the human first originated (Behar et al., 2008; Chan et al., 2015, 2019). In recent years, the researchers found early *H. sapiens* fossils from Morocco which dates far earlier than the other fossils identified till date (Richter et al., 2017; Hublin et al., 2017). These findings raised the debate of the origin of human. Through the genetic study in very recent years, the researchers identified a common ancestor of the populations of the western parts of Africa (Durvasula and Sankararaman, 2020). This ancestor is mentioned as the “ghost archaic human” or “African Neanderthals” (Stringer and Galway-Witham, 2017; Durvasula and Sankararaman, 2020) and carries a significant distinct feature than the Neanderthals and Denisovans. It is also documented that 2 − 19% of the west African DNA are inherited from the ancestor, “African Neanderthals” (Durvasula and Sankararaman, 2020). There are some studies which analyzed the genome-wide data and the artefacts, reported that the ancient humans were more widespread across the globe (Callaway, 2018). Some impressions of the presence of ancient humans earlier than the fossils record added that the ancient human were more widespread than we think (Warren, 2019; Callaway, 2018). In the derived tree, the similar phenomena is also observed, a major clade is formed from the populations of western and central parts of Africa. This clade is placed at the most basal position of the phylogenetic tree with a very high bootstrap support score (see Fig. 3). We estimated the time of origin of the ancestor of the clade as almost 600-700 kya which originated earlier than the other ancient humans, such as Neanderthals and Denisovans, which is further concordance to the opinion of some recent studies (Steinrücken et al., 2018; Schaefer et al., 2016). So this phenomena supports the early occurrence of our species, and this study also supports that the western and central parts of Africa experienced the earliest occurrence of human.

All the derived trees show a very mixed representation of the humans from different communities. Researchers found some signatures of the admixture with modern humans in as recent as 15-30 kya, such that almost 3 − 5% of the Asian DNA are inherited from Denisovans (Gibbons, 2019) and almost 1.5 − 2.1% of the non-African genome are inherited from Neandarthals (Schaefer et al., 2016). Though the bootstrap supports of all of these occurrences are lower than the accepted significant level, but appearance of the same event in a repetitive manner shows a high possibility of the admixture in modern humans.

There are still some small groups placed together in all the derived trees. It is also observed that the Asian populations are closely related to the African population. There are some archaeological studies that report the similarities in the ancient instruments in African and Asian population (Mellars, 2006). Various studies proposed the close relation between north African and Levantine based on the haplogroups of both mtDNA and Y-chromosomes (Olivieri et al., 2006; van de Loosdrecht et al., 2018; Elkamel et al., 2018). It is also noticed that the eastern and the north-eastern Africans are closely placed with both the Asian and the European populations. All the derived trees also show the close relationships among the Pacific islanders (Oceania), such as Micronesia, Melanesia, and Polynesia. It is also observed that DNA of these populations are also close to the DNA of the Taiwanese. This phenomena also supports the existing studies where the close link between the Pacific islanders and the Taiwanese is described (Trejaut, 2005; Friedlaender et al., 2008). Though the bootstrap score is less than that of the accepted level, but its repeated occurrence in the derived trees implies the higher chance of these relationships being true in the phylogenetic study.

All the observations of this study have been summarized in Table 1.

**Table 1:**
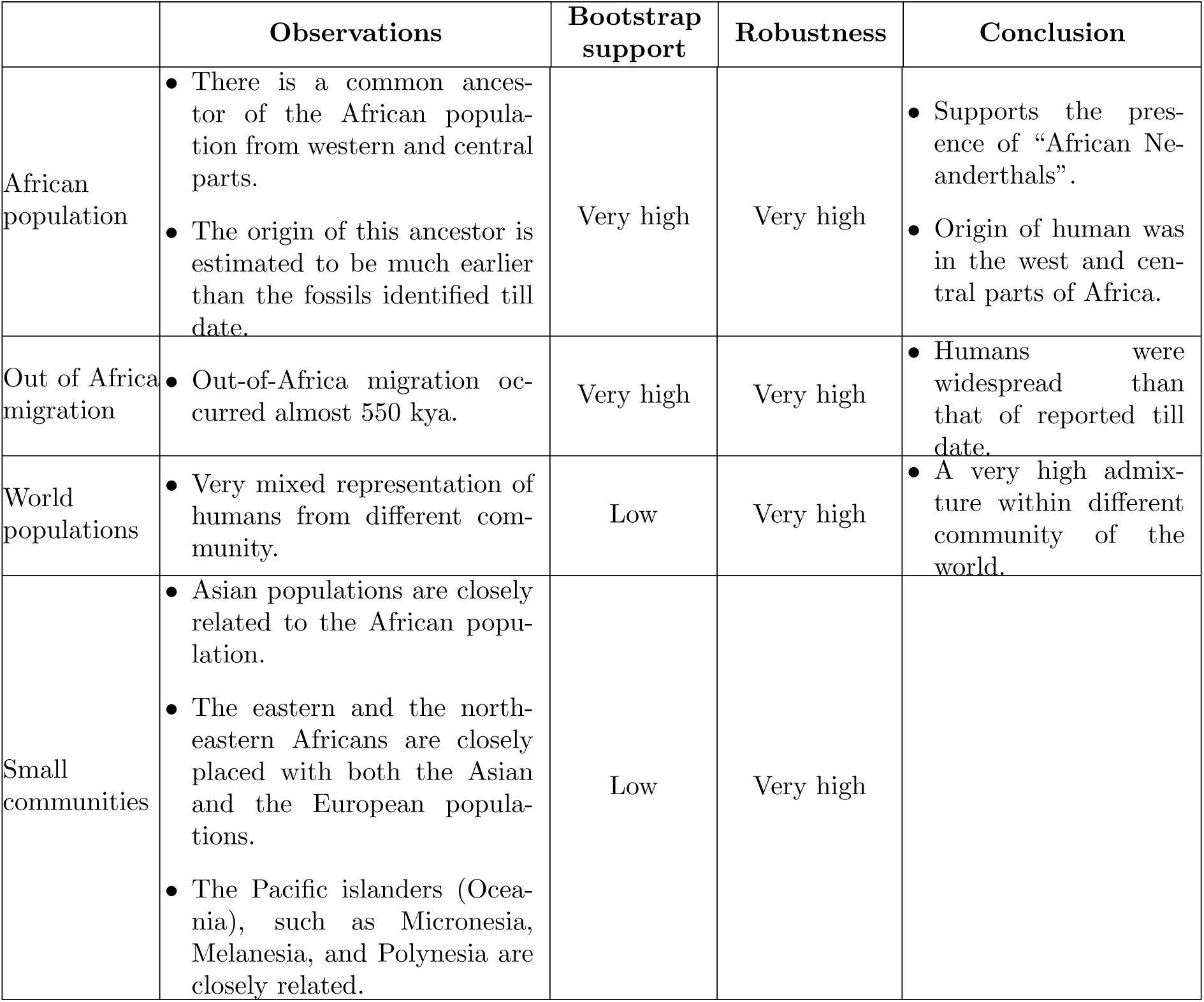
Summary of the observations

## 5 Conclusion

In this study we reported a study of human origin based on the graphical representation where we inspect the origin and the migration of human based on a novel feature vector. The proposed method may be extended to study evolution of other types of genomic sequences. This method captures some unique signature of the whole mtDNA. It is observed that this method efficiently captures the broad relationships rather than the precise relationships near the leaf nodes. In this study, we present such broad divergence among the human population. Our result supports some existing hypotheses like the Africa witnessed the first human in the world and the close relationship among the Eurasian and the African population. The origin of the African clade in our study is also close to the group of archaeological reports. Apart from that, this study provides an evidence of the presence of the “ghost ancestor” of the population of the west and central Africa earlier than that of the fossils identified. This phenomena also demonstrates that the west and central Africa are very important for the early evolution of human.

## Code availability

We implemented a Python based pipeline available at http://www.facweb.iitkgp.ac.in/~jay/GraFree/HIIARG-GRAFree.html. The mirror copy of this program is also available at https://github.com/aritramhp/GRAFree.git. This program (i) computes the tree based on the GRAFree algorithm, (ii) generates multiple number of bootstrap sequences of a target sequence, and (iii) derives the bootstrap trees from the bootstrap sequences.

## Data availability

All the data are publicly available in the online database, NCBI, with their accession number. The list of accession numbers we consider in this study are provided in **Supplementary S1**. The human mitochondrial sequences are further annotated and stored in public database server, mtDB.

## Ethics declarations

### Conflict of interest

On behalf of all authors, the corresponding author states that there is no conflict of interest.

### Ethical Approval

The ethical approval from the institute is not applicable for this study.

### Funding

This research did not receive any specific grant from funding agencies in the public, commercial or not-for-profit sectors.

### Informed Consent

All the data and personal information are collected from a public database. These data have been generated by various researchers since a decade.

## Notes

#### Summary of Updates

1) Rewrite the manuscript for better clarification. 2) Modified the title of the manuscript.

