## Supplementary S2 for "Human origin and migration deciphered from a novel genomic footprints of mitochondrial sequences"

### Supplementary Material for
### "Study of human origin and migration from a novel genomic footprint derived from mitochondiral sequences"

Aritra Mahapatra and Jayanta Mukherjee
*Indian Institute of Technology, Kharagpur, India 721302*

#### 1 Derived trees using GRAFree

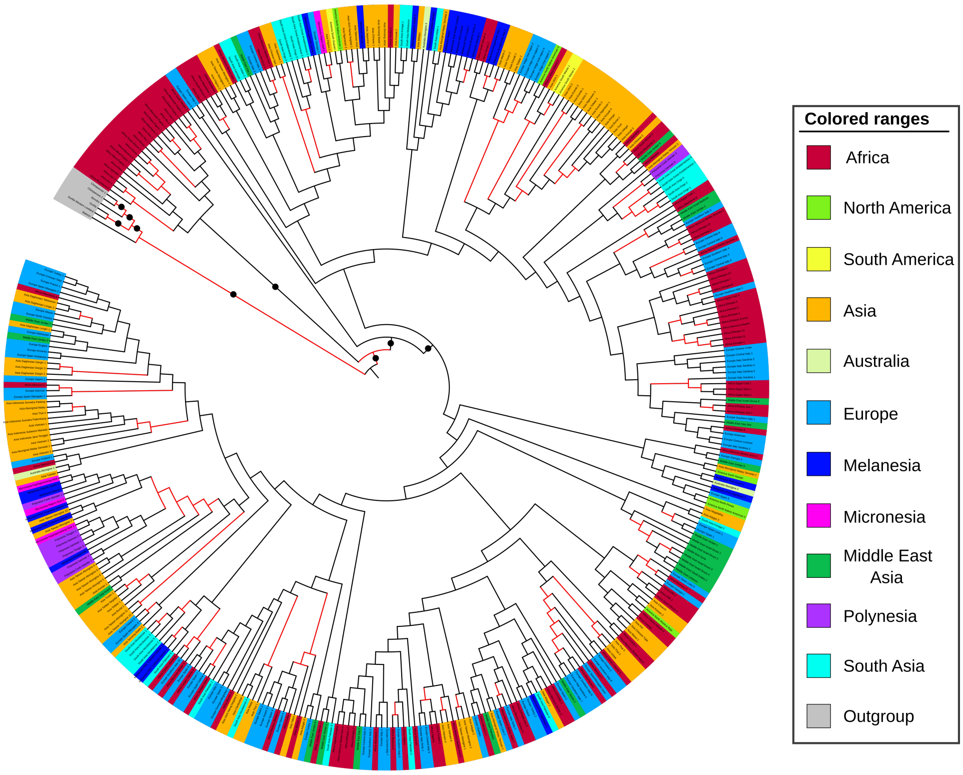


**Figure 1:** Tree is derived for the case when we consider the Purine and Pyrimidine as 2D coordinates space. The black dots on the edge denote a high bootstrap support ($\geq 75\%$) of the corresponding clade. The edge marked as red color denotes that the robustness of the clade with the changes of the hyper parameters is greater than 80%.

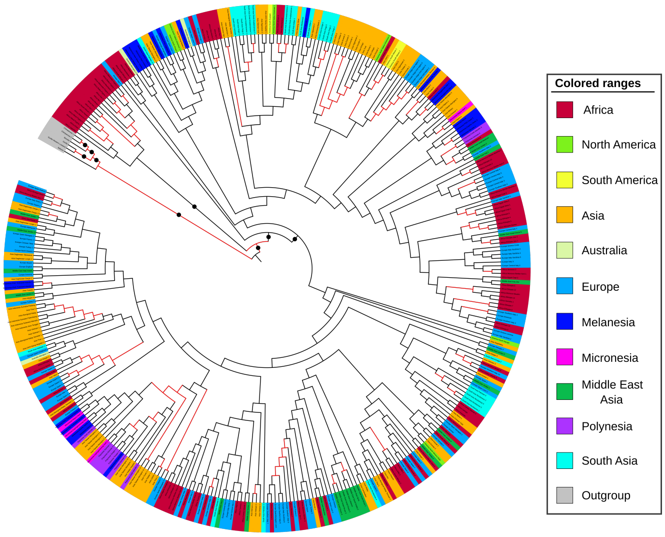


**Figure 2:** Tree is derived for the case when we consider the Strong H-bond and Weak H-bond as 2D coordinates space. The black dots on the edge denote a high bootstrap support ($\geq 75\%$) of the corresponding clade. The edge marked as red color denotes that the robustness of the clade with the changes of the hyper parameters is greater than 80%.

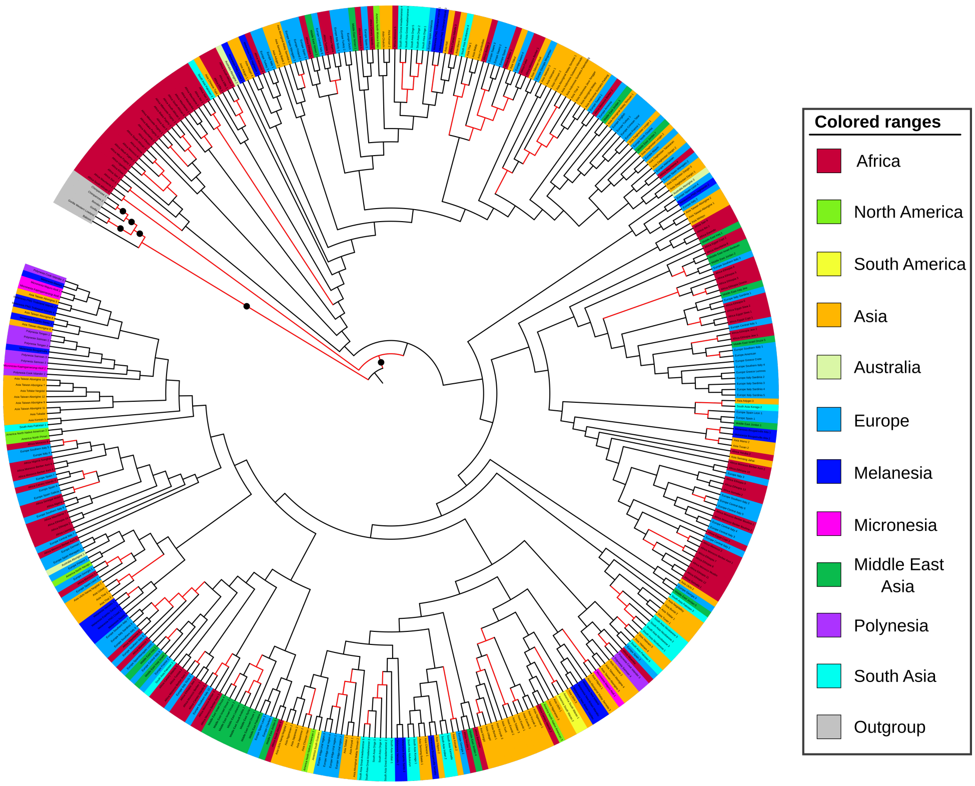


**Figure 3:** Tree is derived for the case when we consider the Amino and Keto as 2D coordinates space. The black dots on the edge denote a high bootstrap support ($\geq 75\%$) of the corresponding clade. The edge marked as red color denotes that the robustness of the clade with the changes of the hyper parameters is greater than 80%.

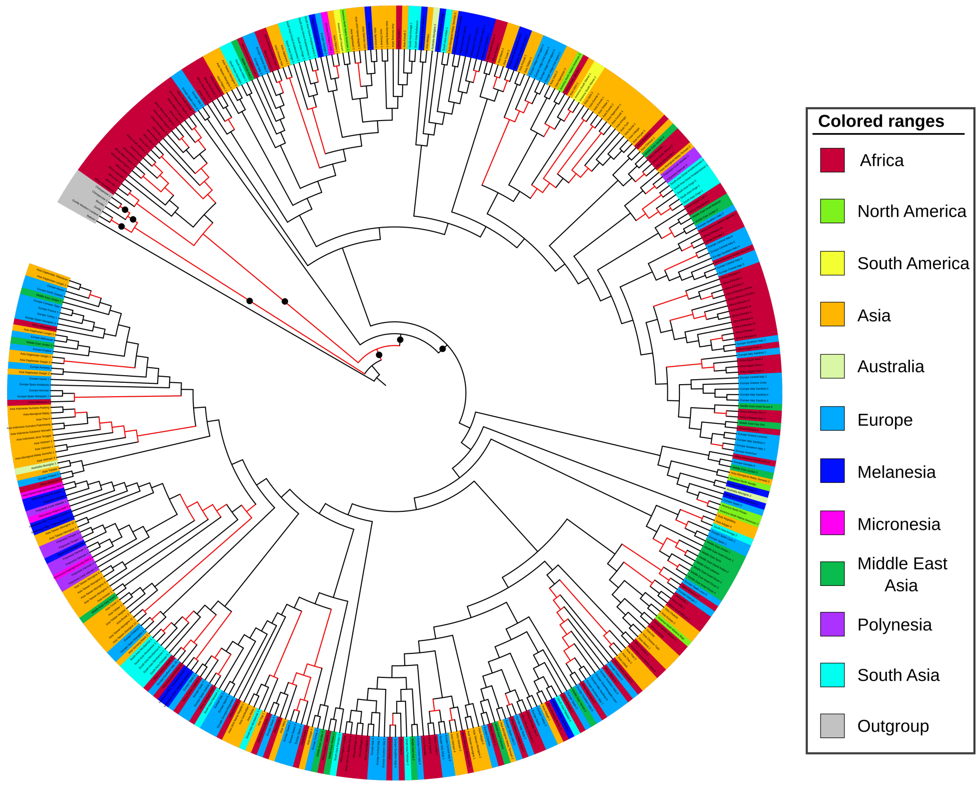


**Figure 4:** Tree is derived after combining three trees derived from three different hypotheses. The black dots on the edge denote a high bootstrap support ($\geq 75\%$) of the corresponding clade. The edge marked as red color denotes that the robustness of the clade with the changes of the hyper parameters is greater than 80%.

#### 2 Derived trees using Euclidean distance function

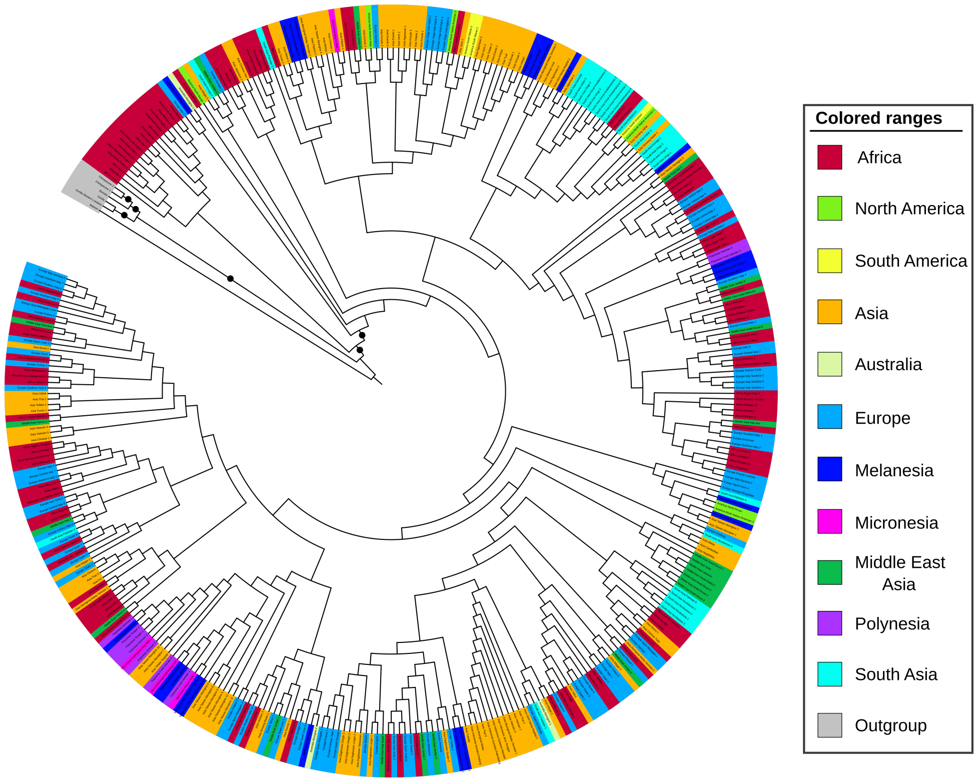


**Figure 5:** Tree is derived after combining three trees derived from three different hypotheses and by considering the Euclidean distance function. The black dots on the edge denote a high bootstrap support ($\geq 75\%$) of the corresponding clade.
